# Developmental regulation of neuronal gene expression by Elongator complex protein 1 dosage

**DOI:** 10.1101/2021.04.16.440179

**Authors:** Elisabetta Morini, Dadi Gao, Emily M. Logan, Monica Salani, Aram J. Krauson, Anil Chekuri, Yei-Tsung Chen, Ashok Ragavendran, Probir Chakravarty, Serkan Erdin, Alexei Stortchevoi, Jesper Q. Svejstrup, Michael E. Talkowski, Susan A. Slaugenhaupt

## Abstract

Elongator is a highly conserved protein complex required for transcriptional elongation, intracellular transport and translation. Elongator complex protein 1 (ELP1) is the scaffolding protein of Elongator and is essential for its assembly and stability. Familial dysautonomia (FD), a hereditary sensory and autonomic neuropathy, is caused by a mutation in *ELP1* that lead to a tissue-specific reduction of ELP1 protein. Our work to generate a phenotypic mouse model for FD led to the discovery that homozygous deletion of the mouse *Elp1* gene leads to embryonic lethality prior to mid-gestation. Given that FD is caused by a reduction, not loss, of ELP1, we generated two new mouse models by introducing different copy numbers of the human FD *ELP1* transgene into the *Elp1* knockout mouse (*Elp1*^*-/-*^) and observed that human *ELP1* expression rescues embryonic development in a dose dependent manner. We then conducted a comprehensive transcriptome analysis in mouse embryos to identify genes and pathways whose expression correlates with the amount of *ELP1*. We found that *ELP1* is essential for the expression of genes responsible for the formation and development of the nervous system. Further, gene length analysis of the differentially expressed genes showed that the loss of *Elp1* mainly impacts the expression of long genes and that by gradually restoring Elongator their expression is progressively rescued. Finally, through evaluation of co-expression modules, we identified gene sets with unique expression patterns that depended on *ELP1* expression. Overall, this study highlights the crucial role of *ELP1* during early embryonic neuronal development and reveals gene networks and biological pathways that are regulated by Elongator.

## Introduction

Elongator is a highly conserved multiprotein complex composed of two copies of each of its six subunits, named Elongator complex proteins 1 to 6 (ELP1-6). Elongator subunits are evolutionarily highly conserved from yeast to humans both in their sequence and interaction with other subunits. Conserved function across all species has been clearly demonstrated using a variety of different cross-species rescue experiments (1-3). Deletion of any of the genes encoding the six subunits confers almost identical biochemical phenotypes in yeast (4-6), suggesting that there is a tight functional association between the proteins comprising Elongator complex, and that the functional integrity of Elongator is compromised in the absence of any of its subunits (6). Both yeast and human Elongator have lysine acetyltransferase activity that is mediated by the catalytic subunit Elp3. Elp3 has two identified substrates: histone H3 and α-Tubulin (7-10). While the acetylation of histone H3 is linked to the role of the complex in transcriptional elongation (7, 10, 11), cytosolic acetylation of α-Tubulin has been linked to its role in microtubule organization particularly in the context of cell migration (8). Elongator was isolated as a complex that associates with chromatin and interacts with the elongating phosphorylated form of RNA polymerase II (RNAPII) both in yeast and human (7, 10, 12). The catalytic subunit Elp3, by acetylating histone H3, facilitates RNAPII access to actively transcribed genes. In human cells, Elongator is required for the expression of several genes involved in migration and in the expression of HSP70 (13, 14). In addition, accumulating evidence supports the role of this complex in maintaining translational fidelity through tRNA modifications. Specifically, Elongator is essential for the formation of 5-methoxy-carbonylmethyl (mcm5) and 5-car-bamoylmethyl (ncm5) groups on uridine nucleosides present at the wobble position of many tRNAs (15, 16).

Several *loss-of-function* studies have demonstrated the key role of Elongator during development. Yeast Elp mutants are hypersensitive to high temperature and osmotic conditions and they showed defects in exocytosis, telomeric gene silencing, DNA damage response and adaption to new growth medium (10, 17, 18). In *Arabidopsis thaliana*, mutations in Elp subunits resulted in impaired root growth (19) and deletion of Elp3 in *Drosophila melanogaster* was lethal at the larval stage (20). Depletion of Elongator in *Caenorhabditis elegans* led to defects in neurodevelopment (21). In mice, Elp1 knockout results in embryonic lethality due to failed neurulation and vascular system formation (3, 22). Consistent with its crucial role during development, several human neurodevelopmental disorders have been associated with mutations in Elongator complex subunits. Familial dysautonomia (FD) is caused by a splicing mutation in *ELP1* (23-25) that reduces the amount of functional protein in a tissue specific manner, missense mutations in *ELP2* have been found in individuals with severe intellectual disability (ID) (26, 27), variants of *ELP3* have been associated with amyotrophic lateral sclerosis (ALS) (28), *ELP4* variants have been implicated in autism spectrum disorder and Rolandic epilepsy (RE) (29, 30) and a mutation in *Elp6* causes Purkinjie neuron degeneration and ataxia-like phenotypes in mice (31).

FD is a neurodevelopmental disorder characterized by widespread sensory and autonomic dysfunction and by central nervous system (CNS) pathology (32-37). The major mutation in FD is a splicing mutation in *ELP1* intron 20 that leads to variable skipping of exon 20 and to a reduction of ELP1 mostly in the nervous system (25, 38). In 2009 we generated a knockout (KO) Elp1 mouse, Elp1^-/-^ and showed that complete ablation of *Elp1* resulted in early embryonic lethality (3, 22). To gain a better understanding how reduction of ELP1 leads to FD, we generated two new mouse models by introducing different copy numbers of the human *ELP1* transgene with the major FD mutation (39), *TgFD1* and *TgFD9*, into the *Elp1*^*-/-*^ mouse. Although the human FD transgene did not rescue embryonic lethality of the *Elp1*^*-/-*^ mouse, its expression rescues embryonic development in a dose dependent manner in Tg*FD1; Elp1*^*-/-*^ and *TgFD9; Elp1*^*-/-*^ embryos. In order to understand the gene regulatory networks that are dependent on ELP1 expression, we conducted a comprehensive transcriptome analysis in these mouse embryos.

## Results

### Generation of mice expressing an increasing amount of *ELP1*

We previously demonstrated that homozygous deletion of the mouse *Elp1* gene leads to embryonic lethality prior to mid-gestation (3). Detailed phenotypic characterization of *Elp1*^*-/-*^ KO embryos at early developmental stages revealed several abnormalities, including a dramatic reduction in size, disruption of the extraembryonic vascular networks, failure of germ layer inversion, and interruption of cephalic neural-tube closure (3, 22). In an effort to understand the molecular mechanisms that characterize FD we have generated several transgenic mouse lines carrying the wild-type (WT) and FD human *ELP1* gene that differ by the copy number of the transgene (39). The murine Elp1 protein is 80% identical to human ELP1 and by introducing the human WT *ELP1* transgene into the *Elp1*^*-/-*^ mouse, we completely rescued development and mice were born alive and healthy, confirming ELP1 functional conservation between human and mouse (3). To test whether the abnormalities caused by ablation of mouse *Elp1* could be improved by the human FD transgene, heterozygote mice carrying different copy numbers of the FD *ELP1* transgene (*TgFD1; Elp1*^*+/-*^ or *TgFD9; Elp1*^*+/-*^) were crossed with mice heterozygous for the *Elp1* knockout allele (*Elp1*^-*/+*^). Mice were collected at either E8.5 or P0 and genotyped using genomic DNA from the visceral yolk sac. Although neither *TgFD1* or *TgFD9* rescued embryonic lethality in the *Elp1*^*-/-*^ mice (Table 1), the development of the FD1/KO (Tg*FD1; Elp1*^*-/-*^) and FD9/KO (*TgFD9; Elp1*^*-/-*^) embryos progressed further as human *ELP1* expression increased (Figure 1a). Because familial dysautonomia is caused by a developmental reduction of ELP1 protein, we conducted a comprehensive transcriptome analysis in embryos expressing increasing amounts of ELP1. We collected 29 individual C57BL/6 mouse embryos at E8.5 (n=8 KO, n=7 FD1/ KO, n=6 FD9/ KO, n=8 WT) and total RNA was extracted from each single embryo (Figure 1a). As expected, KO embryos do not express any WT *Elp1* (Figure 1b); whereas FD1/KO and FD9/KO embryos expressed increasing amounts of WT *ELP1* with FD9/KO embryos expressing three times more *ELP1* than FD1/KO embryos (Figure 1c). Normalized gene count comparisons revealed that the expression of the human *ELP1* in FD1/KO embryos is ∼6% of WT *Elp1* while in the FD9/KO the human *ELP1* is ∼16% WT *Elp1*. Principal component analysis revealed that the four genotypes exhibited distinct expression profiles with PC1 explaining 28% of the total expression variance across samples suggesting that *ELP1* dosage plays a critical role in embryonic transcriptome regulation (Figure 1d).

**Table 1.** Genotype ratios of offspring generated by *TgFD; Elp1*^*+/-*^*X Elp1*^*+/-*^ pairings. The time of conception is estimated to be E0.5 day prior to the observation of a vaginal plug. P0, postnatal day zero. Expected mendelian ratio is indicated below the genotype, percentage of embryos of each genotype in parentheses.

| *TgFD; Elp1*<sup>+/-</sup><br>X<br>*Elp1*<sup>+/-</sup> | Time | Number of animals |  |  |  |  |  | Total |
| --- | --- | --- | --- | --- | --- | --- | --- | --- |
|  |  | *Elp1*<sup>+/+</sup><br>12.5% | *Elp1*<sup>+/-</sup><br>25% | *Elp1*<sup>-/-</sup><br>12.5% | *Tg; Elp1*<sup>+/+</sup><br>12.5% | *Tg; Elp1*<sup>+/-</sup><br>25% | *Tg; Elp1*<sup>-/-</sup><br>12.5% |  |
| *TgFD1* | E8.5 | 17(11%) | 37(25%) | 19(13%) | 17(11%) | 43(29%) | 17(11%) | 150 |
| *TgFD1* | P0 | 43(19%) | 88(39%) | 0(0%) | 29(13%) | 63(28%) | 0(0%) | 223 |
| *TgFD9* | E8.5 | 10(7%) | 30(21%) | 16(11%) | 25(17%) | 45(31%) | 17(11%) | 143 |
| *TgFD9* | P0 | 24(23%) | 42(40%) | 0(0%) | 13(12%) | 27(25%) | 0(0%) | 106 |

**Figure 1.**
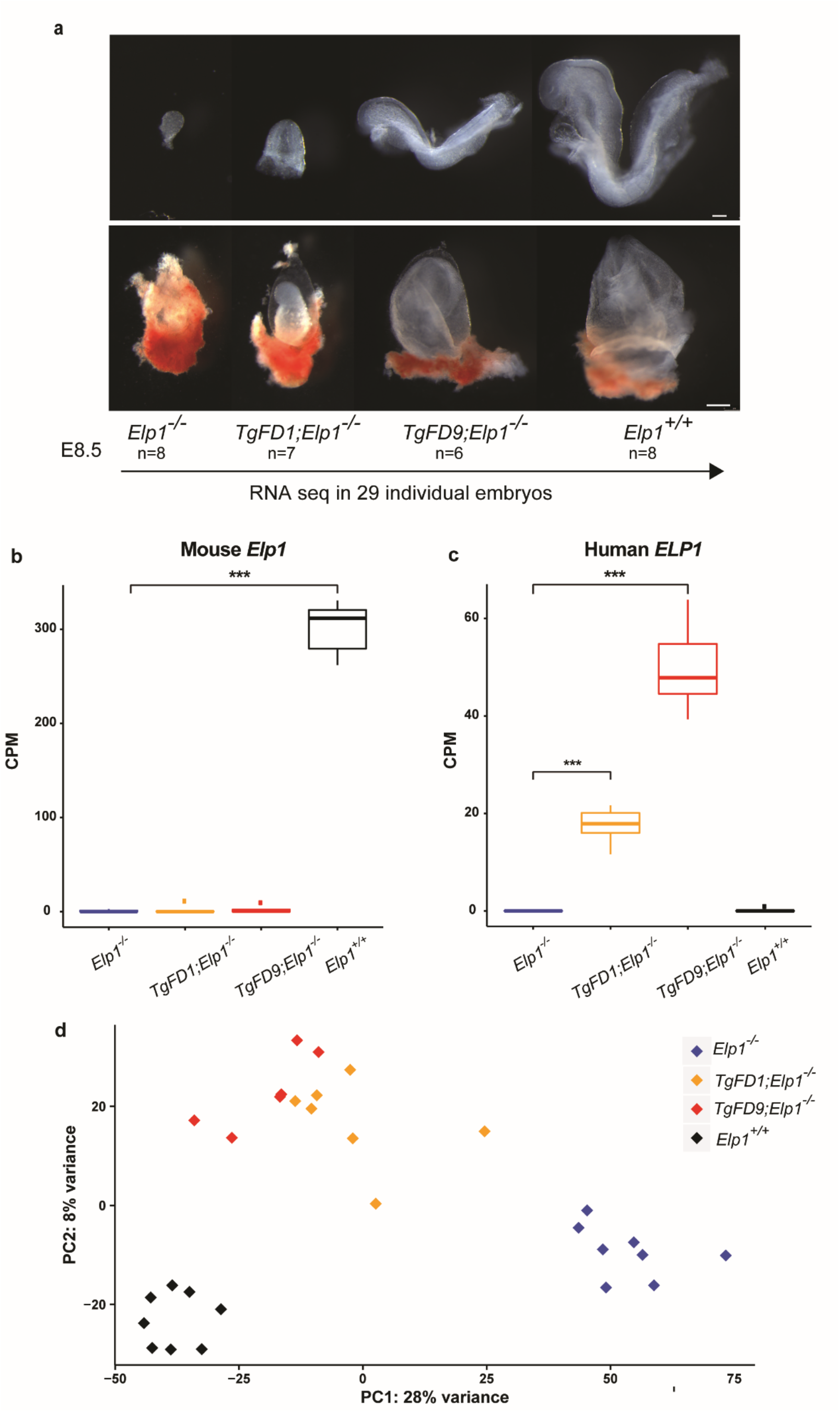
Generating mouse embryos expressing increasing amount of ELP1. (**a**) Morphological phenotypes of *Elp1*^*-/-*^, *TgFD1; Elp1*^*-/-*^, *TgFD9; Elp1*^*-/-*^ and *Elp1*^*+/+*^ embryos (top, the scale bar represents 100 µm) and extraembryonic components (bottom, the scale bar represents 250 µm) at E8.5. RNA-seq experiment was performed using total RNA extracted by individual embryo. (**b**) Expression of the endogenous WT *Elp1* across different embryos. The median for each group is shown. (**c**) Expression of the human WT *ELP1* across different embryos. The median for each group is shown. (**d**) Principal component analysis of all the embryos colored by genotype. In box-and-whisker plots in **b**-**c**, each box extends to 1.5 times inter-quartile range (IQR) from upper and lower hinges, respectively. Outliers are not shown. Only comparisons with significant difference are marked by stars (two-tailed, unpaired Welch’s *t* test with Bonferroni correction). In the figure, \**P* < 0.05; \*\**P* < 0.01; \*\*\**P* < 0.001.

### Major transcriptome changes in *Elp1* KO embryos

Transcriptome profiling in embryos expressing an increasing amount of *ELP1* show that the number of differentially expressed genes (DEGs), (False Discovery Rate or FDR < 0.1; and Fold Change or FC >= 1.5), proportionally declines as *ELP1* expression increases. In KO embryos 2,399 out of 19,619 (12.23%) genes were differentially expressed when compared with WT embryos (Figure 2a, Supplementary Table 1). Strikingly, in FD1/KO embryos the DEGs were only 601 (3.06%), while in FD9/KO embryos there were 494 DEGs (2.52%) (Figure 2a, Supplementary Table 1), demonstrating that a minimal increase in *ELP1* is sufficient to rescue the expression of ∼80% of all DEGs. Gene Ontology (GO) analysis of the down-regulated genes (FDR < 0.1) in KO, FD1/KO and FD9/KO embryos highlighted pathways important for nervous system development including synapse formation, neuron projection and axon growth (Figure 2b, Supplementary Table 2). These findings are consistent with the body of work supporting the role of *ELP1* during early development in target tissue innervation and with the fact that neuronal loss in FD is mostly due to failure of innervation (14, 40-45). Notably, of the 71 genes that were significantly downregulated in all three KO genotypes (Supplementary Table 1), 24 of them (∼33%) have a critical role in nervous system or brain development (Figure 2c). STRING analysis of these genes revealed an enrichment for protein-protein interactions (PPI enrichment = 2.9E-4 according to STRING v11) (Figure 2c) (46). Among the neuronal genes, *Dbx1* and *Nr2e1* were the two most down-regulated genes across all three KO genotypes (Figure 2d and 2e). *Dbx1*, also known as Developing Brain Homeobox 1, is expressed in a regionally restricted pattern in the developing mouse central nervous system (CNS) and encodes for a transcription factor that plays a pivotal role in interneuron differentiation in the ventral spinal cord (47). In vertebrates, spinal interneurons modulate the motor neuron activity elicited by incoming sensory information and, by relaying the proprioceptive data to the brain, play a critical role in locomotor coordination (48). *Nr2e1* is a transcription factor that regulates the expression of genes essential for retinal development (49). Loss of *Nr2e1* in mice has been shown to cause severe early onset retinal degeneration with death of various retinal cells including retinal ganglion cells (RGCs) (50-52). Interestingly, the list of neuronal genes that were significantly downregulated in all three KO genotypes, KO, FD1/KO and FD9/KO, also included the brain-derived neurotrophic factor BDNF-receptor TrkB (*Ntrk2)*, the chemorepulsive axon guidance protein draxin, the homeobox protein involved in brain and sensory organ development otx2, and the neuronal adhesion protein involved in neurite growth neurocan (*Ncan*) (Figure 2f-i). GO pathways associated with up-regulated genes in the KO embryos are included in Supplementary Figure S1and Supplementary Table 2.

**Figure 2.**
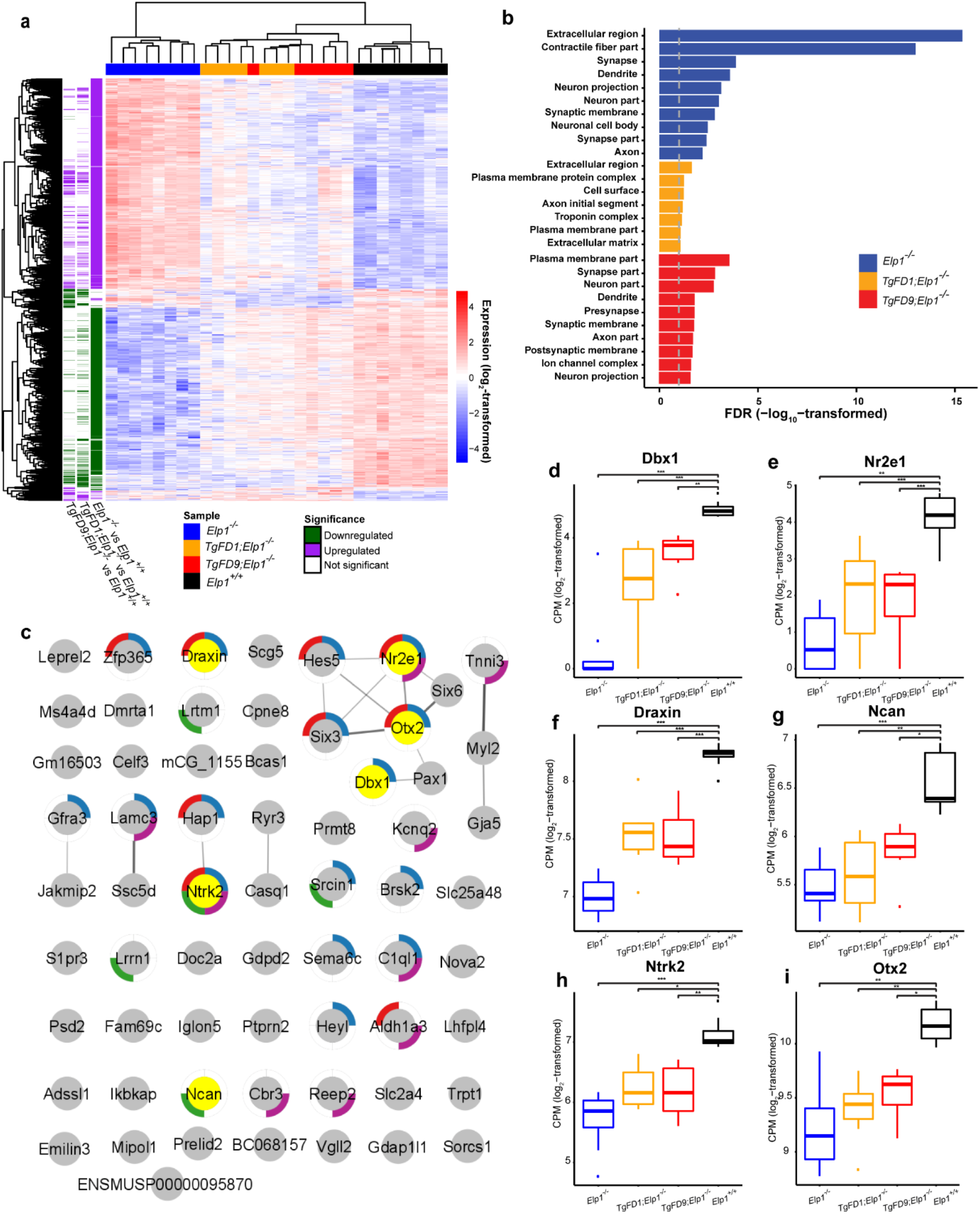
Transcriptome profiling of embryos expressing increasing amount of ELP1. (**a**) Heatmap of the 2,619 differentially expressed genes for each genotype compared to WT *Elp1*^*+/+*^. (**b**) Gene ontology analysis of the downregulated genes in *Elp1*^*-/-*^, *TgFD1; Elp1*^*-/-*^, *TgFD9; Elp1*^*-/-*^embryos. The graph shows FDR values for the most significant specific GO terms (see also Supplementary Table 2). (**c**) STRING interaction map for the 71 genes that were significantly downregulated in all three KO genotypes (see also Supplementary Table 1). Each pie indicates a gene. Note, *Ikbkap* is the alternative name of *Elp1*. The pies surrounded by colored edges indicate genes involved in neurogenesis (GO:0022008, *blue*), brain development (GO:0007420, *red*), regulation of synapse organization (GO:0050807, *green*), and nervous system process (GO:0050877, *purple*), respectively. The pies in yellow indicate the genes whose expression are highlighted in panels d-i. (**d**-**i**) box-and-whisker plots showing the expression of key genes for nervous system development that are significantly downregulated across all three KO genotypes: Dbx1 (**d**), Nr2e1 (**e**), Draxin (**f**), Ncan (**g**), Ntrk2 (**h**) and Otx2 (**i**). \**P* <0.05, \*\**P* <0.01 and \*\*\**P* < 0.001, Welch’s *t* test.

### Long neuronal genes require Elongator activity for their expression

Although Elongator plays a number of roles in the cell, in the nucleus this complex directly interacts with RNAPII and facilitates transcriptional elongation through altering chromatin structure (7, 10, 12, 53). Elongator has histone acetyltransferase activity via its ELP3 subunit and regulates the accessibility of RNAPII to the chromatin. Using Chromatin immunoprecipitation (ChIP) we have previously shown that in the absence of ELP1, histone H3 acetylation was significantly reduced at the 3’ ends of genes (3). Moreover, Close et al demonstrated that upon reduction of ELP1 there was progressively lower RNAPII density at the 3’ end of target genes than in the promoter region (14), supporting the role of Elongator in transcriptional elongation. To examine if the downregulation in gene expression observed in the KO embryos might be due to a failure in transcriptional elongation, we compared the length distribution of the DEGs among embryos expressing an increasing amount of *ELP1*. We discovered that long genes, especially those longer than 100 kb, were downregulated significantly more often in the KO embryos than in the FD9/KO embryos (FDR < 0.01, Figure 3a) suggesting that Elongator loss has a more pronounced effect on the expression of longer genes. Importantly, ELP1 restoration efficiently rescued their expression. In contrast, there was no difference in gene length in the upregulated genes between the different genotypes (Figure 3b). Of the 257 long genes (> 100 kb) that were downregulated in KO embryos, 216 (84.05%) were rescued in the FD1/KO embryos and 247 (96.11%) were rescued in FD9/KO embryos. GO analysis of these genes highlighted pathways important for synapse formation, neuron projection and axon growth (Figure 3c, Supplementary Table 3). Since long downregulated genes were enriched for pathways important for nervous system development, we compared the average length of neurodevelopmental genes with the average length of all expressed genes (see Material and Methods) and we observed that neurodevelopmental genes (GO:0048666) were significantly longer (Figure 3d). We then investigated whether the ELP1-dependent gene regulation was driven by length or if neuronal genes were more likely to require ELP1 for efficient transcription. We divided all downregulated genes in KO embryos into four gene-length categories and examined the proportion of neurodevelopmental genes (GO:0048666) in each category (Figure 3e *left panel*). If neurodevelopmental genes were more likely to require ELP1 for efficient transcription, we would expect to see more rescue with higher ELP1 dosage. Given that the proportion of neurodevelopmental genes rescued in the FD9/KO embryos in all gene-length categories was similar to those of the downregulated genes in the KO embryos (Figure 3e *right panel*), we concluded that neurodevelopmental genes are more susceptible to ELP1-loss simply because they are longer than the non-neurodevelopmental genes.

**Figure 3.**
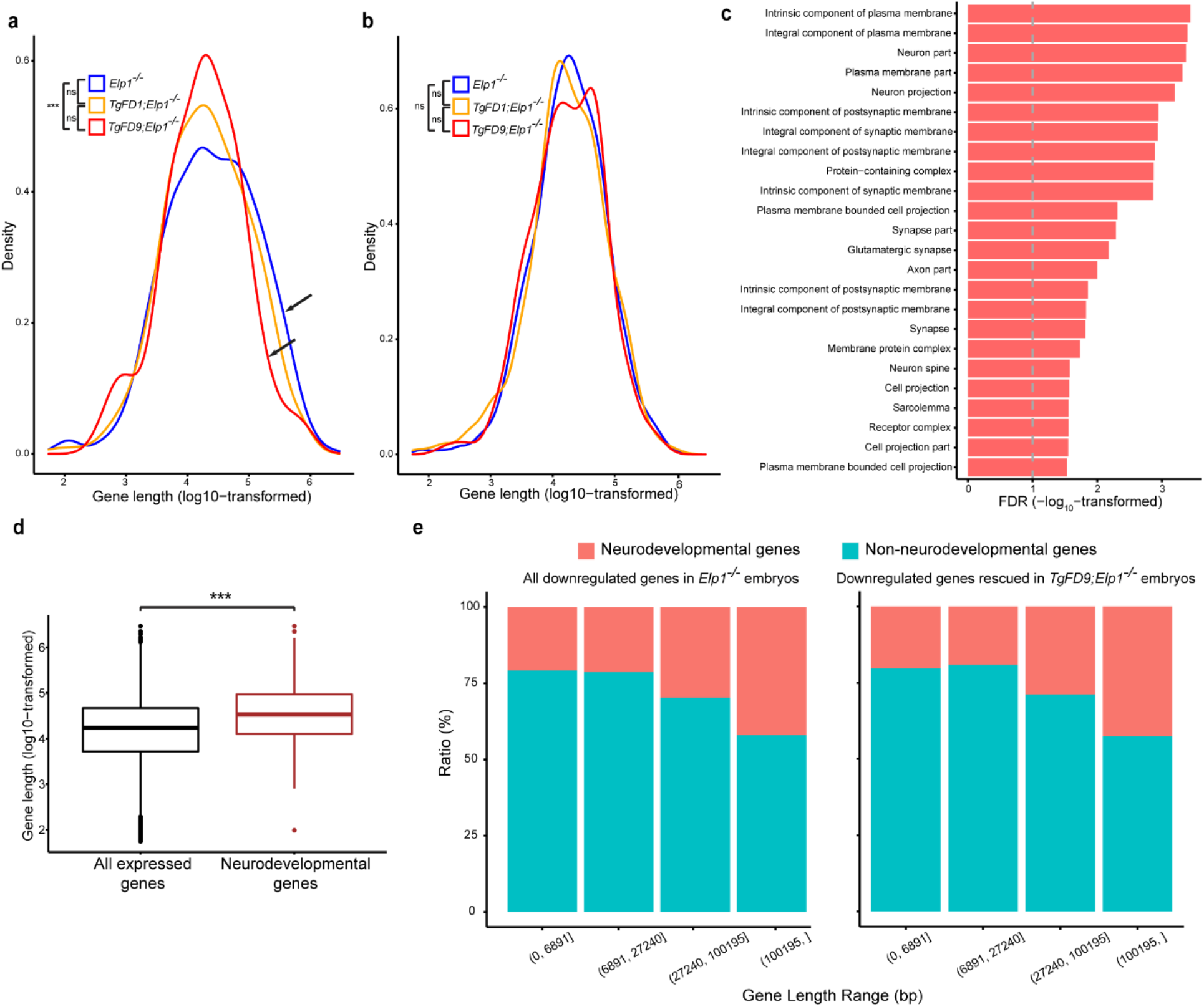
Gene length distribution in embryos expressing different ELP1 amounts. (**a**) Length distribution of the down-regulated genes in *Elp1*^*-/-*^, *TgFD1; Elp1*^*-/-*^ and *TgFD9; Elp1*^*-/-*^ embryos. Arrows indicate the comparison between *Elp1*^*-/-*^ and *TgFD9; Elp1*^*-/-*^ distribution. (**b**) Length distribution of the up-regulated genes in *Elp1*^*-/-*^, *TgFD1; Elp1*^*-/-*^ and *TgFD9; Elp1*^*-/-*^ embryos. In both **(a)** and **(b)**, K-S test was applied. (**c**) Gene ontology analysis of the 247 long genes (> 100 kb) downregulated in *Elp1*^*-/-*^ embryos and rescued at *TgFD9; Elp1*^*-/-*^. The graph shows FDR values for each specific GO term (see also Supplementary Table 3). **(d)** Comparison between length distributions of all expressed genes in RNASeq with the length distribution of neuronal genes (GO:0048666 in Gene Ontology database) via two-tailed, unpaired Welch’s *t* test. **(e)** Ratio of neurodevelopmental and non-neurodevelopmental genes in different gene-length ranges. The *left* panel represents all downregulated genes in *Elp1*^*-/-*^ embryos, while the *right* panel represents genes which expression was rescued in *TgFD9; Elp1*^*-/-*^ embryos. The gene-length ranges were determined in order to have an equal number of genes in each range. In the figure, ns, not significant and \**P* < 0.05; \*\**P* < 0.01; \*\*\**P* < 0.001.

### Identification of genes whose expression correlates with amount of ELP1

Elongator has been linked to transcriptional regulation (13, 14, 54). Our unique mouse models provide, for the first time, the ability to perform a comprehensive transcriptome analysis to identify genes whose expression depends on the amount of *ELP1*. This study is highly relevant to better understanding FD pathogenesis, as the disease is caused by a reduction, not loss, of *ELP1* primarily in the nervous system. We built the gene co-expression network across embryonic RNASeq data for all genotypes (Materials and Methods). We identified thirty-five co-expression distinct Modules Eigengenes (MEs) (Figure 4a). We then postulated that genes whose expression relies on *ELP1* expression would be grouped into three major categories (Figure 4b): (1) genes whose expression changes as a monotonic function of *ELP1*, referred as “dose-responsive genes”; (2) genes whose expression is completely rescued with low *ELP1* expression, referred as “highly responsive genes” and (3) genes whose expression is restored only when *ELP1* is expressed at WT levels, referred as “low responsive genes”. Among the thirty-five MEs identified, ME3, ME2 and M12 had the highest positive correlation with these hypothesized gene patterns (Pearson correlation >= 0.85, FDR < 0.1, Figure 4c). ME3 included the dose-responsive genes (Figure 4d); ME2 included the highly responsive genes (Figure 4g) and M12 contained the low responsive genes (Figure 4j). Interestingly, ME4, ME7 and ME10 had the highest negative correlation with our hypothesized patterns (Supplementary Figure S2).

**Figure 4.**
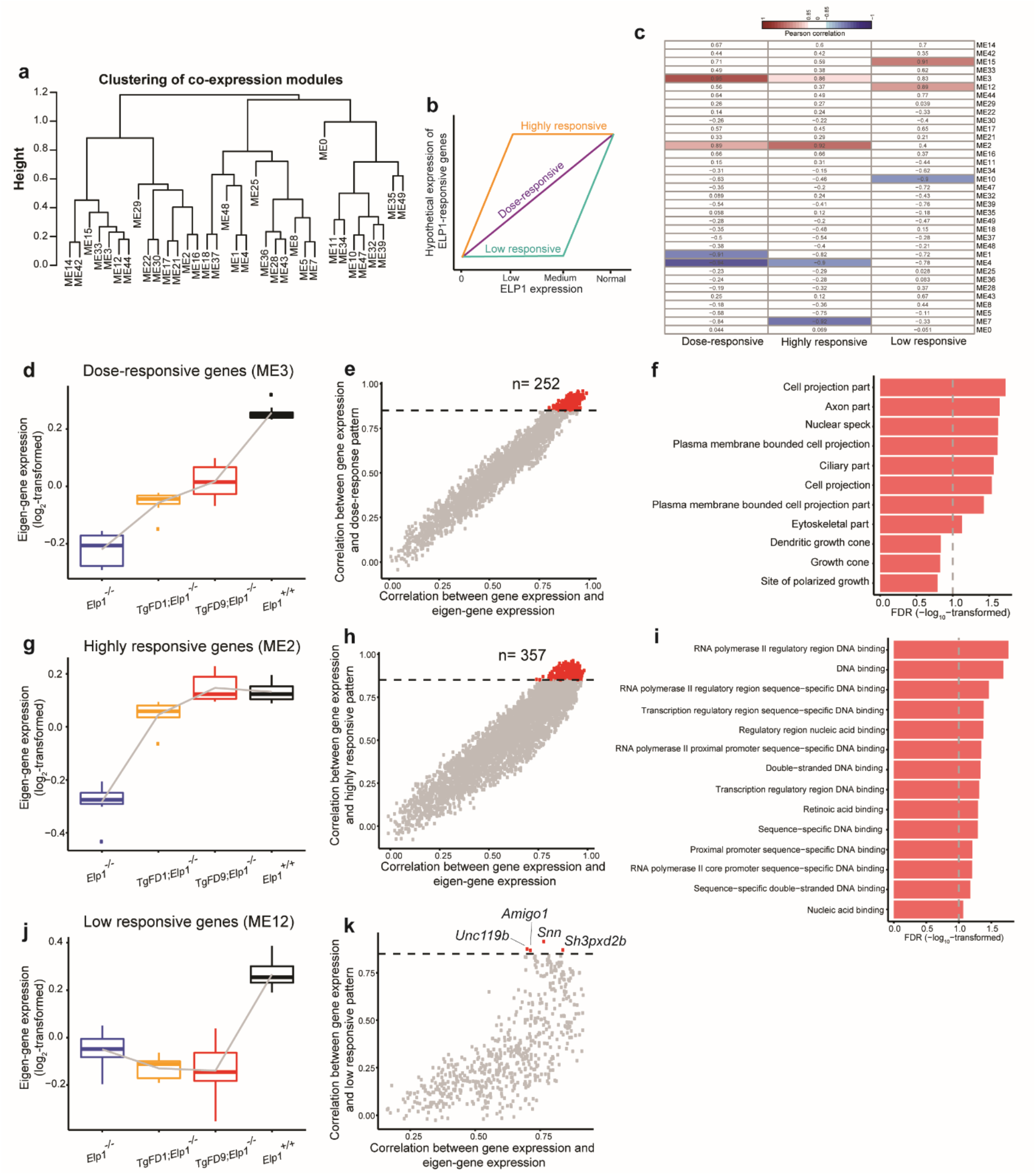
Identification of *ELP1*-responsive gene patterns. (**a**) Modules of distinct Eigen (ME) analysis across embryos expressing increasing amount of *ELP1* identified thirty-five distinct MEs (see also Materials and Methods). (**b**) Hypothetical expression trajectories of *ELP1*-responsive genes. (**c**) Heatmap displays the Pearson correlation between the eigen values of each co-expression module and the eigen values of each hypothetical expression trajectory. The blue domain indicates a negative correlation while the red domain indicates a positive correlation. The number in each grid demonstrates the Pearson correlation coefficient. (**d**) Boxplot displays the eigen values of gene expression of ME3 at each genotype. (**e**) Module membership of each gene in ME3. Each dot represents a gene. The x-axis demonstrates the Pearson correlation between gene expression and module eigen values of ME3 while the y-axis demonstrates the Pearson correlation between gene expression and the eigen values of each hypothetical expression trajectory. The horizontal dashed line shows a correlation coefficient of 0.85. (**f**) Gene ontology analysis of the 252 dose-responsive genes which expression strictly increase as monotonic function of *ELP1*. The graph shows FDR values for each specific GO term (see also Supplementary Table 4). (**g**) Boxplot displays the eigen values of gene expression of ME2 at each genotype. (**h**) Module membership of each gene in ME2. (**i**) Gene ontology analysis of the 357 highly responsive genes whose expression is completely restored in *FD9/KO* embryos. The graph shows FDR values for each specific GO term (see also Supplementary Table 4). (**j**) Boxplot displays the eigen values of gene expression of ME12 at each genotype. (**k**) Module membership of each gene in ME12. \**P* < 0.05; \*\**P* < 0.01; \*\*\**P* < 0.001, Welch’s *t* test.

Overall, we identified 242 dose-responsive genes whose expression strictly increases as a monotonic function of *ELP1* (Pearson correlation >= 0.85, FDR < 0.1, Figure 4e). The GO analysis of these genes highlighted pathways important for axon and cell projection formation (Figure 4f, Supplementary Table 4). This result supports, once again, the role of ELP1 in the expression of genes important for target tissue innervation and is consistent with the innervation failure observed in FD (14, 40-45). We found 357 highly responsive genes whose expression was completely restored in *FD9/KO* embryos (Pearson correlation >= 0.85, FDR < 0.1, Figure 4h). GO analysis of these genes highlighted pathways associated with transcriptional regulation (Figure 4i, Supplementary Table 4) suggesting that a small increase of functional Elongator is enough to restore normal expression of important transcriptional regulators. This finding might explain the dramatic phenotypic improvement observed in *FD9/KO* embryos when compared with *FD1/KO* and KO. Finally, we identified four low responsive genes whose expression was rescued solely in WT embryos: *Amigo1, Snn, Unc119b* and *Sh3pxd2b* (Figure 4k) (Pearson correlation >= 0.85, FDR < 0.1).

In the negatively correlated patterns ME4, ME7 and ME10 (Pearson correlation >= 0.85, FDR < 0.1) we did not find any enrichment for neuronal terms (Supplementary Figure S2, Supplementary Table 5). Genes in ME4 highlighted pathways involved in mitochondrial and respiratory chain activity (Supplementary Figure S3, Supplementary Table 5) while genes in ME7 and ME10 did not shown any significant enrichment for GO terms (FDR < 0.1). These results show that the expression of many genes that are involved in transcriptional regulation and nervous system development positively correlate with *ELP1* expression.

## Discussion

Variants in Elongator subunits are associated with various human neurodevelopmental disorders, including FD (*ELP1*), ID (*ELP2*), ALS (*ELP3*), autism spectrum disorder and Rolandic epilepsy (*ELP4*) (23-31). To gain a better understanding the pathogenesis of FD, we generated two new mouse models expressing an increasing amount of *ELP1*, FD1/KO and FD9/KO. Although the human FD transgene did not rescue embryonic lethality in the *ELP1* KO mouse, its expression improved embryonic development in a dose dependent manner. To identify genes and pathways whose expression is highly correlated with *ELP1* and that are ultimately essential for embryonic development, we conducted a comprehensive transcriptome analysis in KO, FD1/KO, FD9/KO and WT embryos. We found that even a minimal increase in *ELP1* has a dramatic effect on overall gene expression with the majority of the KO DEGs being completely rescued in the FD1/KO embryos, which only expresses an amount of *ELP1* that is approximately 6% of the *Elp1* amount expressed in the WT embryos. A significant portion of the down-regulated genes across the different genotypes KO, FD1/KO and FD9/KO, have a crucial role in nervous system development. Among these neuronal genes, *Dbx1* and *Nr2e1* were the two most down-regulated genes in all three genotypes with absent or reduced expression of *ELP1*.

*Dbx1*, also known as *Developing Brain Homeobox 1*, is expressed in a regionally restricted pattern in the developing mouse central nervous system (CNS) and encodes for a transcription factor that has pivotal role in interneuron differentiation in the ventral spinal cord (47). The spatially restricted expression of *Dbx1* is important to establish the distinction of V0 neurons from the adjacent set of V1 neurons (55). *Dbx1* mutant mice lacking these ventral interneurons exhibit marked changes in motor coordination, supporting the role of Dbx1-dependent interneurons as key components of the spinal locomotor circuits that control stepping movements in mammals (48). Interestingly, one of the most characteristic features of FD is poor locomotor coordination and both patients as well as a phenotypic mouse model of FD showed progressive impairment in coordination that leads to severe gait ataxia (56-58).

*Nr2e1*, or nuclear receptor subfamily 2 group E member 1, encodes a highly conserved transcription factor known to be a key stem cell fate determinant in both the developing mouse forebrain and retina (52, 59-63). The mouse *Nr2e1* gene is first detected at embryonic day 8 (E8) in the ventricular zone (VZ) of the neuroepithelium layer and later spreads posteriorly into the diencephalon (E10.5) (63-66). In the developing mouse eye, *Nr2e1* is detected in the optic processes of the developing embryo as early as E9 (32), suggesting a role for this gene in an early phase of retinogenesis (29, 61). During development, *Nr2e1* null mice display an increase in apoptotic levels of RGCs in the ganglion cell layer (GCL), which results in a marked reduction in thickness of the distinct layers in the adult retina and optic nerve dystrophy (51, 52, 66). Intriguingly, degeneration of RGCs is observed in two different *Elp1* conditional knock-out mice (67, 68). In addition, patients with FD show RGCs loss with reduction in the thickness of the retinal nerve fiber layer (RNFL) and progressive vision loss (33, 34, 69). Although further studies will be necessary to determine the link between *ELP1* reduction and downregulation of specific key neurodevelopmental genes, the identified *ELP1*-dependent transcriptome profiles constitute an excellent foundational resource for understanding Elongator biology and also help to shed light into the molecular pathways that underlie diseases caused by disruption of Elongator activity. In the nucleus, Elongator facilitates transcriptional elongation through altering chromatin structure (3, 7, 10, 53). Therefore, we analyzed the length distribution of DEGs among embryos expressing different *ELP1* amounts. Our data clearly showed that long genes are more affected by the loss of functional Elongator compared with shorter genes and, by gradually increasing *ELP1* amount, we were able to progressively restore their expression. Moreover, the observation that neuronal genes are significantly longer than all expressed genes, offers a possible explanation about why the nervous system is the tissue that most relies on functional Elongator during embryonic development. In the future it would be interesting to investigate the role of *ELP1* in transcriptional elongation for each identified target, in order to identify direct regulatory effects.

Several genes that require Elongator to be efficiently expressed have been identified using either cell lines or conditional KO mouse tissues with reduced *Elp1* expression (14, 45, 70-73). In addition, RNA microarray analysis in post-mortem FD tissues has shown that a subset of genes involved in myelination require *ELP1* for efficient transcription (74). In the current study we have identified gene patterns whose expression varies as a function of *ELP1* amount: (1) genes whose expression changes as a monotonic function of *ELP1*; (2) genes whose expression is completely restored with low amount of *ELP1* and (3) genes whose expression is restored only when *ELP1* is expressed at WT levels (Figure 4c). It is interesting that distinctive groups of genes display different Elongator dependence. Genes important for axon formation and cell projection responded in a dose-dependent manner to functional Elongator amounts. This is consistent with the observation that, even though Elongator is a ubiquitously expressed protein complex, variants affecting different Elongator subunits all lead to neurodevelopmental diseases (23-31). On the other hand, the expression of important transcriptional regulators is already restored in *FD9/KO* embryos, suggesting that only a small increase of functional Elongator is necessary to rescue their expression. This might underlie the dramatic phenotypic improvement observed in *FD9/KO* embryos when compared with *FD1/KO* and KO. We then identified sets of genes whose expression negatively correlated with *ELP1*. GO analysis of these genes highlighted pathways involved in mitochondrial and respiratory chain activity.

In conclusion, our study is the first to assess the *in vivo* dose-dependent effect of *ELP1* in early development using transcriptome analysis. We demonstrated that even a minimal increase in *ELP1* can have a dramatic effect on both mouse embryonic development and global gene expression. Although loss of *ELP1* compromised the expression of a large number of genes, this study shows that neuronal genes are more sensitive to *ELP1* reduction. We recognize that further studies will be necessary to determine which of the individual gene expression changes are direct versus indirect. However, the identification of gene networks and biological pathways that are regulated by *ELP1*-dosage is highly relevant to better understand the pathogenesis of FD, as well as other neurodevelopmental diseases caused by Elongator deficiency. The data presented here will help to identify potential biomarkers for future clinical studies and new targetable pathways for therapy.

## Materials and Methods

### Generation of *FD1/KO, FD9/KO* mouse models and genotyping

The detailed description of the original strategy to generate the *Elp1* knockout mouse line has been previously published (3). A detailed description of the generation the *TgFD* transgenic lines carrying different copy number of the human *ELP1* gene with the IVS20+6T>C mutation can be found in our previous manuscript by Hims et al. (39). To create the *TgFD1; Elp1*^*-/-*^ and *TgFD9; Elp1*^*-/-*^ mouse, we crossed the previously generated *TgFD1* or *TgFD9* transgenic mouse line with the mouse line heterozygous for the null allele *Elp1*^*-/+*^. The resulting progeny was genotyped to detect the presence of the *TgFD1* or *TgFD9* transgene and of the null allele *Elp1*^*-*^. As expected, the *TgFD1* and *TgFD9* transgene segregated independently from the null allele; therefore, around one-fourth of the F1 mice carried both the *TgFD* transgene and null *Elp1* alleles (*TgFD1; Elp1*^*+/-*^ or *TgFD9; Elp1*^*+/-*^). Subsequently, we crossed the *TgFD1; Elp1*^*+/-*^ and *TgFD9; Elp1*^*+/-*^ mice with the mouse line heterozygous for the null Elp1 allele (*Elp1*^*+/-*^). The resulting progeny was genotyped to detect the presence of the TgFD1 or TgFD9 transgene as well as the null allele in homozygosis (*Elp1*^*-/-*^). The *Elp1*^*-/-*^, *TgFD1; Elp1*^*-/-*^, *TgFD9; Elp1*^*-/-*^ and *Elp1*^*+/+*^ embryos were produced by crossing heterozygote mice carrying different copy numbers of the FD *ELP1* transgene (*TgFD1; Elp1*^*+/-*^ or *TgFD9; Elp1*^*+/-*^) with heterozygote mice (*Elp1*^-*/+*^). The day of vaginal-plug discovery was designated E 0.5. The mice used for this study were housed in the animal facility of Massachusetts General Hospital (Boston, MA), provided with constant access to a standard diet of food and water, maintained on a 12-hour light/dark cycle, and all experimental protocols were approved by the Subcommittee on Research Animal Care at the Massachusetts General Hospital. The genotypes of animals and embryos were determined by PCR analysis of genomic DNA from tail biopses and from embryos and/or visceral yolk sacs, respectively. The primer sets used were as follows: for determining the wild-type Elp1 allele, 5’-ACCCTCAGGCAGTTTGATTG-3’ and 5’-CATGGCTCCATAAAACAAACAC-3’; for detecting the knockout allele, 5’ACCCTCAGGCAGTTTGATTG-3’ and 5’-GGCTACCGGCTAAAACTTGA-3’; and for determining the human TgFD transgenes, TgProbe1F 5’-GCCATTGTACTGTTTGCGACT-3’ and TgProbe1R 5’-TGAGTGTCACGATTCTTTCTGC-3’.

### Morphological analysis of embryos

Photographs of visceral yolk sacs and embryos were taken with a digital camera Leica DFC7000 T mounted on a Leica M205 FCA dissection microscope. LAS X software (Leica) was used for image processing.

### RNA Seq experiment

RNA was extracted from 8 *Elp1*^*-/-*^, 7 *TgFD1; Elp1*^*-/-*^, 6 *TgFD9; Elp1*^*-/-*^ and 8 *Elp1*^*+/+*^ individual embryos at the embryonic stage of E8.5 using the QIAzol Reagent and following the manufacturer’s instructions. RNAseq Libraries were prepared using TruSeq® Stranded mRNA Library Prep kit (Illumina 20020594), using 100 ng total RNA as input. Libraries were evaluated for final concentration and size distribution by Agilent 2200 TapeStation and/or qPCR, using Library Quantification Kit (KK4854, Kapa Biosystems), and multiplexed by pooling equimolar amounts of each library prior to sequencing. Pooled libraries were 50 base pair paired-end sequenced on Illumina HiSeq 2500 across multiple lanes. Real time image analysis and base calling were performed on the HiSeq 2500 instrument using HiSeq Sequencing Control Software (HCS) and FASTQ files demultiplexed using CASAVA software version 1.8.

A synthesized transcriptome reference was generated, by artificially adding the sequence of human *ELP1* gene from the Ensembl human transcriptome reference GRCh37.75 to the Ensembl mouse transcriptome reference GRCm38.83 as an independent chromosome. RNASeq reads were mapped to this synthesized transcriptome reference 3 by STAR v2.5.2b, allowing only uniquely mapped reads with 5% mismatch (75).

### Differential gene expression analysis

Gene counts were generated by HTSeq-count v0.11.2 with “-s reverse” option to be compatible with Illumina TruSeq reads, according to the gene annotations of the synthesized transcriptome reference. Four genotypes were defined as “FD0”, “FD1”, “FD9” and WT to reflect the amounts of human and mouse ELP1. Gene counts across samples were filtered so that only genes whose median expression amounts were no less than 0.1 counts-per-million-reads in at least one genotype were kept for the downstream analysis. The R (v3.6.1) package “SVA” v3.32.1 was implemented on the filtered gene expression matrix to estimate surrogated variables (SVs) among samples. A generalized linear model was built by the R package “DESeq2” v1.24.0 to correlated gene expression to genotypes reflecting human *ELP1* and mouse *Elp1* amounts, together with all the estimated SVs. During the differential gene expression analysis, the FC and FDR of each genotype was estimated per gene. PCA analysis was implemented based on the top 500 most variable genes across samples, from the filtered gene expression matrix with SVs corrected.

### GO analysis

GO analysis was done by GOrilla (76) (http://cbl-gorilla.cs.technion.ac.il/). Organism was set as *Mus musculus*. Two unranked lists of genes were used for each GO analysis. The GO analysis for DEGs used the following two lists: 1) the list of DEGs for a genotype and 2) the list of all genes in the filtered gene expression matrix. The GO analysis for co-expression modules used the following two lists: 1) the list of genes highly correlated with a hypothetical pattern in a co-expression module and 2) the list of all genes in the filtered gene expression matrix. In the figures related to the GO analysis for DEGs, ten significant terms in the ontology of Cellular Component were selected to be plotted in Figure 2b. The complete list of DEG GO terms for each genotype can be found in Supplementary Table 2.

### Mouse neuronal genes

Mouse neuronal genes were extracted and downloaded via AmiGO2 from geneontology.org, using the key word ‘neuron’ and restricting organism as ‘Mus musculus’. Totally 3,442 unique neuronal genes were found.

### Co-expression modules analysis

R package “WGCNA” (77) v1.68 was implemented to the filtered gene expression matrix with SVs corrected. The soft-thresholding power was determined to be 5. The minimal module size was set to 30. The raw modules were merged using a dis-similarity cutoff of 0.15.

### Correlation between co-expression modules and hypothetical patterns

Dummy expression data were generated to mimic three hypothetical patterns, namely the monotonic increase pattern, the monotonic decrease pattern, and the saturated pattern. The eigen vector representing each co-expression module was then correlated with each of the hypothetical patterns using Pearson correlation.

### Statistical Analysis

All raw p values in this study, if multiple tests were involved, were corrected by the Benjamini-Hochberg Procedure and converted to FDR values. Wald test was applied in the differential gene expression analysis. A significant DEG of a genotype, compared to WT, was defined as FDR < 0.1 and the FC >= 1.5 for that genotype. Fisher’s exact test was used for GO analysis, where a significant enrichment for a GO term was defined as FDR < 0.1. Kolmogorov-Smirnov test (K-S test) was applied to compare the gene length distribution of genes from different groups. A significant difference in length distribution between two groups was defined as FDR < 0.1. A significant correlation throughout this study was defined as the Pearson correlation coefficient >= 0.85 and FDR < 0.1. For all box plots, the middle lines inside boxes indicated the medians. The lower and upper hinges corresponded to the first and third quartiles. Each box extended to 1.5 times inter-quartile range (IQR) from upper and lower hinges respectively. The symbols *, ** and ***, if appeared in the figures, indicated FDR < 0.1, < 0.01 and < 0.001, respectively.

## Acknoledgments

We thank Dr. Lucy Norcliffe-Kaufmann and Dr. Horacio Kaufmann of the Dysautonomia Treatment and Evaluation Center at New York University Medical School for their long-standing collaboration and helpful discussions. We are also grateful to Dr. Frances Lefcort for her comments on the manuscript. This work was supported by National Institutes of Health (NIH) grants (R37NS095640 to S.A.S.), the Francis Crick Institute (Cancer Research UK FC001166 to PC and JQS), the UK Medical Research Council (FC001166 to PC and JQS) and the Welcome Trust (FC001166 to PC and JQS).

