## Supplementary Figures for "Developmental regulation of neuronal gene expression by Elongator complex protein 1 dosage"

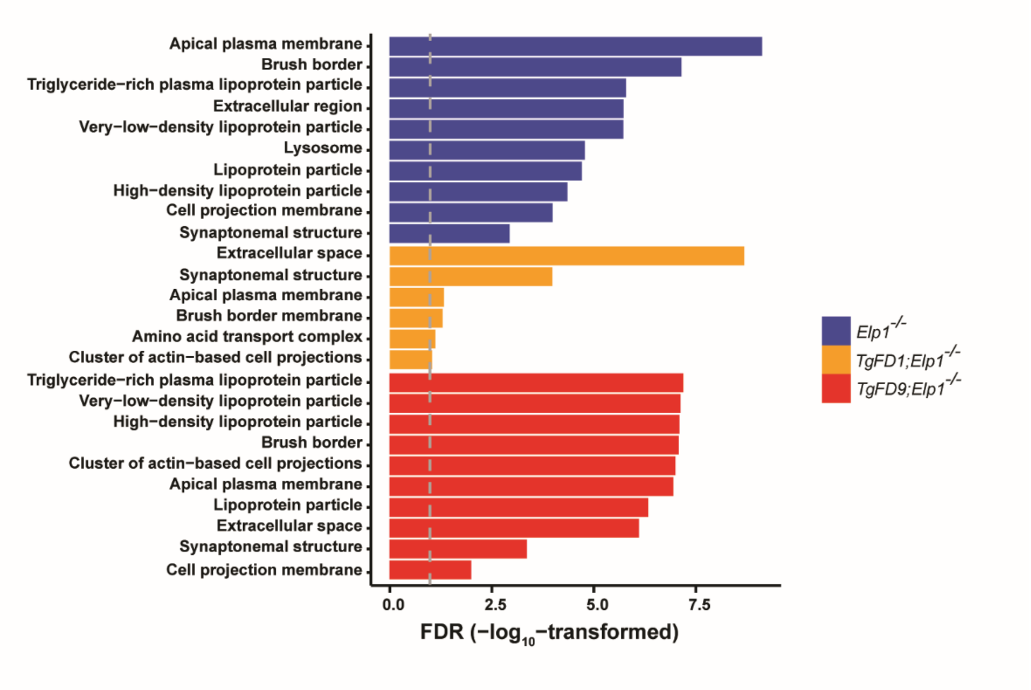


**Supplementary Figure 1. Gene ontology analysis of the upregulated genes in *Elp1$^{-/-}$*, *TgFD1; Elp1$^{-/-}$* and *TgFD9; Elp1$^{-/-}$* embryos.** The graph shows FDR values for the most significant specific GO terms (see also Supplementary Table 2).

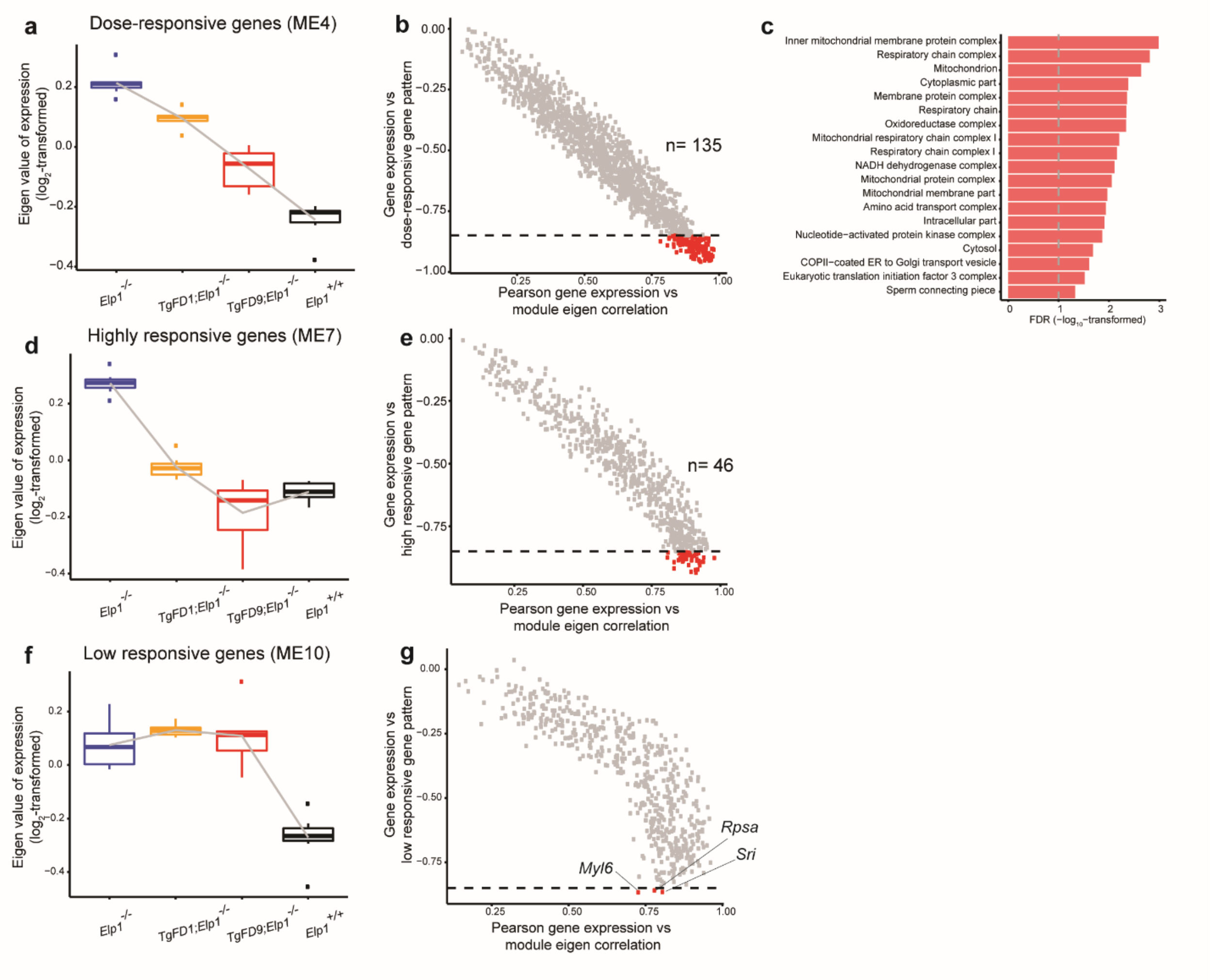


**Supplementary Figure 2. Responsive gene patterns that negatively correlate with *ELP1* amounts.** (**a**) Boxplot displays the eigen values of gene expression of ME4 at each genotype. (**b**) Module membership of each gene in ME4. Each dot represents a gene. The x-axis demonstrates the Pearson correlation between gene expression and module eigen values of ME4 while the y-axis demonstrates the Pearson correlation between gene expression and the eigen values of each hypothetical expression trajectory. The horizontal dashed line shows a correlation coefficient of 0.85. (**c**) Gene ontology analysis of the 135 dose-responsive genes which expression strictly decrease as monotonic function of ELP1. The graph shows FDR values for each specific GO term (see also Supplementary Table 4). (**d**) Boxplot displays the eigen values of gene expression of

ME7 at each genotype. (**e**) Module membership of each gene in ME7. (**f**) Boxplot displays the eigen values of gene expression of ME10 at each genotype. (**g**) Module membership of each gene in ME10. *$P < 0.05$; **$P < 0.01$; ***$P < 0.001$, Welch's *t* test.
